# Reconstructing Neuronal Circuitry from Parallel Spike Trains

**DOI:** 10.1101/334078

**Authors:** Ryota Kobayashi, Shuhei Kurita, Katsunori Kitano, Kenji Mizuseki, Barry J. Richmond, Shigeru Shinomoto

**Author notes:** Electronic mail (Shigeru Shinomoto).

## Abstract

State-of-the-art techniques allow researchers to record large numbers of spike trains parallel for many hours. With enough such data, we should be able to infer the connectivity among neurons. Here we develop a computationally realizable method for reconstructing neuronal circuitry by applying a generalized linear model (GLM) to spike crosscorrelations. Our method estimates interneuronal connections in units of postsynaptic potentials and the amount of spike recording needed for verifying connections. The performance of inference is optimized by counting the estimation errors using synthetic data from a network of Hodgkin-Huxley type neurons. By applying our method to rat hippocampal data, we show that the numbers and types of connections estimated from our calculations match the results inferred from other physiological cues. Our method provides the means to build a circuit diagram from recorded spike trains, thereby providing a basis for elucidating the differences in information processing in different brain regions.

## I. INTRODUCTION

Over the past decade it has become possible to record from much larger numbers of neurons than in the past (Wilson and McNaughton, 1993; Nicolelis *et al.*, 2003; Smith and Kohn, 2008; Schwarz *et al.*, 2014; Mitz *et al.*, 2017), even though this number is still a mere shadow of the total number of neurons present. The premise behind collecting these large data sets is that it will lead to improvements in correlating neuronal activity with specific sensations, motion, or memory, and possibly including their adaptation and learning (Brown, Kass, and Mitra, 2004; Hatsopoulos, Joshi, and O’Leary, 2004; Pillow *et al.*, 2008; Ohiorhenuan *et al.*, 2010; Stevenson and Kording, 2011).

Having such large data sets leads to difficulties in handling the data and interpreting the results. There are two approaches to handling the data. In the first, researchers have developed methods to reduce dimensionality while minimizing the loss of information (Yu *et al.*, 2009; Churchland *et al.*, 2012; Cunningham and Byron, 2014; Williamson, Sahani, and Pillow, 2015; Kobak *et al.*, 2016).

The second approach, which we take here, is to use all of the data to carry out mesoscopic neuroanatomy, that is, to reveal the fine neuronal circuitry in which neural circuit computation is carried out. From the high channel count recordings, one should be able to estimate interneuronal connectivity by quantifying the degree to which firing from a given neuron is influenced by the firing of neurons from which the index neuron is receiving input (Perkel, Gerstein, and Moore, 1967; Toyama, Kimura, and Tanaka, 1981; Reid and Alonso, 1995; Okatan, Wilson, and Brown, 2005; Fujisawa *et al.*, 2008; Stevenson *et al.*, 2009; Chen *et al.*, 2011; Kim *et al.*, 2011; Mishchencko, Vogelstein, and Paninski, 2011; Ito *et al.*, 2011; Stetter *et al.*, 2012; Kobayashi and Kitano, 2013; Zaytsev, Morrison, and Deger, 2015; Gerhard, Deger, and Truccolo, 2017). For this purpose, we develop an analytical tool that estimates interneuronal connectivity in measurement units of postsynaptic potentials (PSPs). In this study we also investigate how much data is needed to reliably estimate the connections between pairs of neurons; that is, we attempt to use data to reconstruct the connectional map among the recorded neurons. The method is evaluated for its accuracy in estimating connections using synthetic data generated by simulating a network of Hodgkin-Huxley (HH)-type neurons. Finally, we apply this method to spike trains recorded from rat hippocampus. For the experimental data, we compare our estimates of whether an innervating connection is excitatory or inhibitory with the results obtained by manually analyzing other physiological information such as spike waveforms, autocorrelograms, and mean firing rate.

## II. RESULTS

### A. Estimating interneuronal connections

To estimate interneuronal connectivity between each pair of neurons, we obtain the cross-correlation (CC) by collecting spike times of a neuron measured relative to every spike of a reference neuron (Figure 1A). We explore the CC for evidence of a mono-synaptic impact of a few milliseconds using the generalized linear model (GLM). Here, interneuronal connectivity is detected by fitting a coupling filter, while slow, large scale wavy fluctuations that are often present in recorded spike trains are absorbed by adapting the slow part of the GLM (METHODS).

**FIG. 1.**
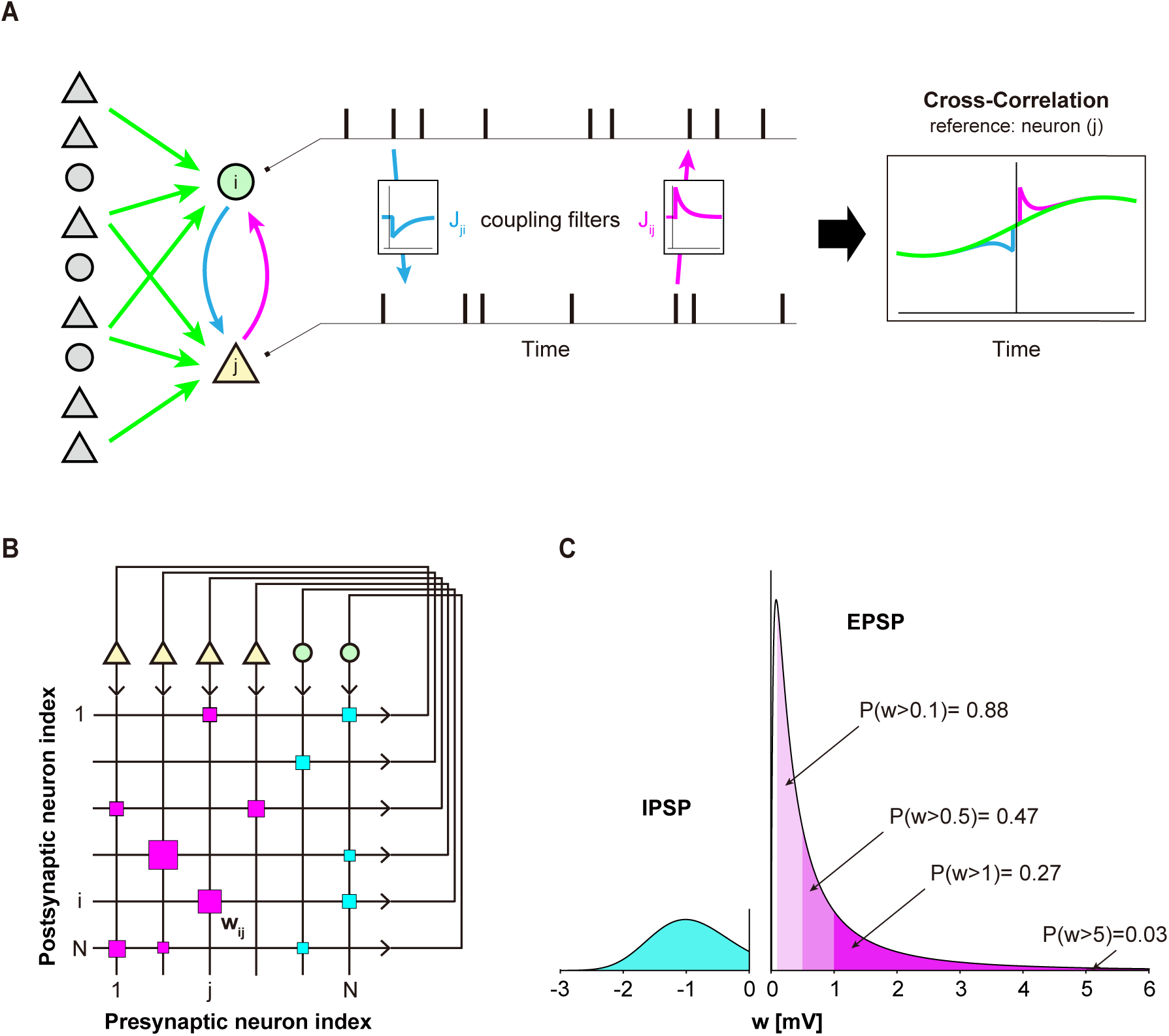
Estimating interneuronal connections. (A) Connectivity between neurons is estimated by fitting a generalized linear model (GLM) to the cross-correlation (CC). *J*_*ij*_ represents a coupling from the *j*-th neuron to the *i*-th neuron. Excitatory and inhibitory neurons are depicted as triangles and circles, and their synaptic connections are colored magenta and cyan, respectively. Surrounding neurons may induce large-scale fluctuations in the CC (green). (B) Interneuronal connectivity is visualized by the Hinton diagram, in which excitatory and inhibitory connections are represented, respectively by magenta and cyan squares of the sizes (area) proportional to the postsynaptic potential (PSP) *w*_*i j*_. (C) Distributions of excitatory postsynaptic potentials (EPSPs) and inhibitory postsynaptic potentials (IPSPs) of a simulated network.

### B. Criterion for the presence of connections

An interneuronal connection is considered significant when the estimated parameter falls outside the confidence interval of a given *p* value for the null hypothesis that the connection is absent. If the parameter remains within the confidence interval, the state of the connection is undetermined (METHODS). The number of pairs considered to be connected will depend on the *p* value and on the strength of the correlation. Our method estimates connections as if they were all direct, making all but certain that strong indirect influences will be converted into purported direct connections. Neurophysiologists often try to avoid these so-called false positives by shifting the *p* value to small values, that is, moving *p* to very stringent levels. However, being conservative about false positives means that existing connections important for information processing will be missed, thereby producing many false negatives.

To examine the influences of false positives and false negatives reflect the level of conservatism used for estimating connections, we applied our inference model to spike trains obtained from a network of HH neurons, in which the true anatomical connectivity is known. We studied at what values the level of conservatism for the *p* value seemed to balance the conflicting demands optimally.

Our simulation used a network of 1,000 HH neurons consisting of 800 excitatory and 200 inhibitory neurons (Figure 1B). In the simulation, excitatory neurons innervated 12.5% of other neurons with excitatory postsynaptic potentials (EPSPs). These excitatory connections were log-normally distributed (Song *et al.*, 2005; Teramae, Tsubo, and Fukai, 2012; Ikegaya *et al.*, 2013; Buzsáki and Mizuseki, 2014) (Figure 1C). Inhibitory neurons innervated 25% of other neurons with inhibitory postsynaptic potentials (IPSPs) randomly. These inhibitory connections were normally (Gaussian) distributed (Hoffmann *et al.*, 2015). We simulated the network for a period representing 5,400 s (90 min) with step sizes of 0.01 and 0.001 ms for excitatory and inhibitory neurons, respectively (METHODS).

To illustrate the performance of estimating connections, we sampled 20 neurons out of the entire population. Figure 2A shows the estimated connection matrices obtained using different *p* values. The connection matrix is divided into four quadrants representing connections between inhibitory-excitatory, excitatory-excitatory, excitatory-inhibitory, and inhibitory-inhibitory neurons. True connections for the 2nd and 3rd quadrants are excitatory, and those of the 4th and 1st quadrants are inhibitory. For *p* = 0.01, too many false connections were assigned to pairs of neurons; there were 15 false connections or false positives out of 318 unconnected pairs (4.7%) in this sample. At the other extreme, all false positives can be excluded by decreasing the *p* value (down to *p* = 10^*−*24^). In the latter case most existing connections are lost, and a large number of false negatives arise; 59 among 62 existing connections (95%) are missed in this example. The numbers of false positives and false negatives for excitatory and inhibitory categories are shown below for the connection matrices, indicating that the total number of false positives and false negatives may be minimized between these extreme cases.

**FIG. 2.**
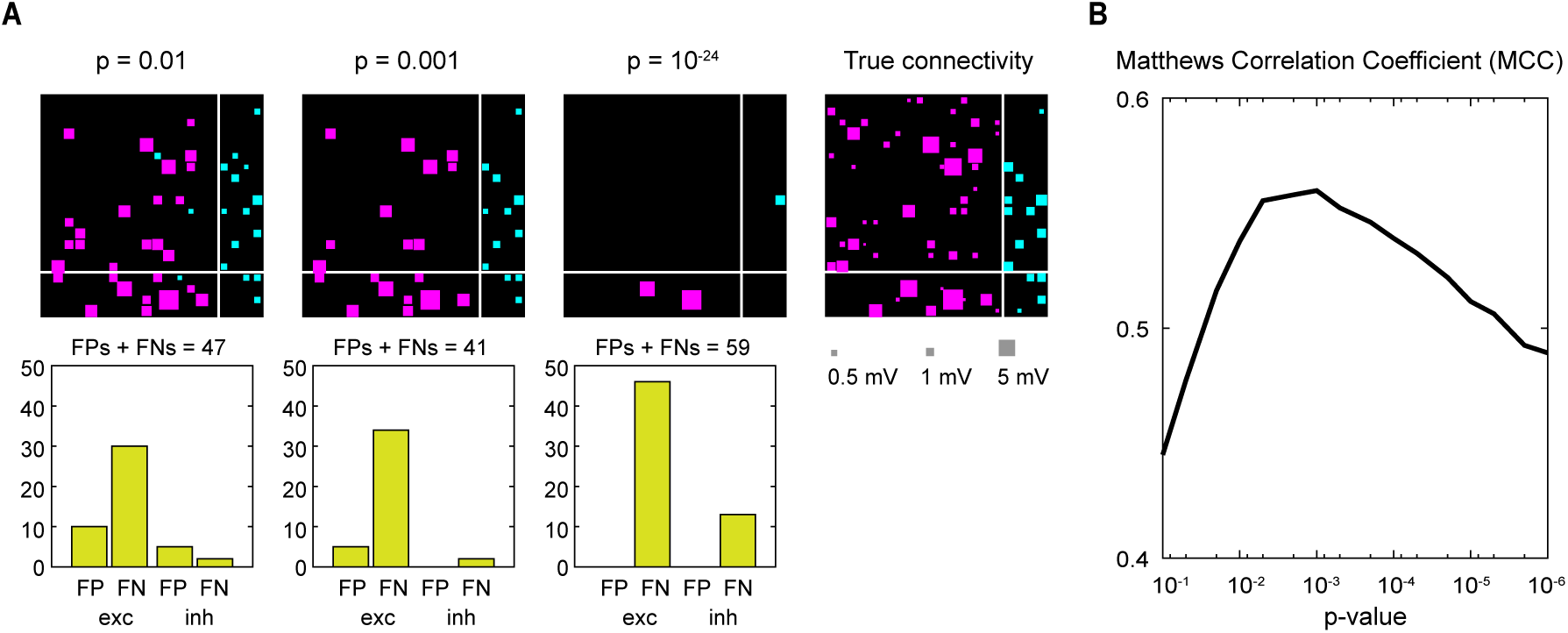
Selecting the *p* value. (A) The connection matrices are estimated with different levels of conservatism against making false positives (FPs) represented by the *p* values. In each connection matrix, *x*-axis indicates reference (index) neurons. The connection matrix is divided into four quadrants, representing inhibitory-excitatory, excitatory-excitatory, excitatory-inhibitory, and inhibitory-inhibitory zones. The numbers of FP and false negative (FN) connections for excitatory and inhibitory categories are depicted below the matrices. (B) The Matthews correlation coefficient (MCC) is plotted against the *p* value. The MCC takes a maximum at an intermediate level of cautiousness given by *p* = 0.001.

To balance the false positives and false negatives simultaneously, we selected the *p* value that maximized the Matthews correlation coefficient (Matthews, 1975; Kobayashi and Kitano, 2013). The *p* value was set to *p* =0.001 (Figure 2B). Although false connections remain, the neuronal circuit was most accurately reconstructed with *p* = 0.001. We adopted *p* = 0.001 throughout the following analyses.

### C. Duration of spike recording

The necessary duration of spike recording can be estimated even without fitting the statistical model to the spike trains. This is because the distribution of the connection parameter for the null hypothesis is obtained solely in terms of the observation interval (*T*) and the firing rates of the preand postsynaptic neurons (*λ*_pre_ and *λ*_post_) (METHODS). The confidence interval of the connection parameter (*J*) is

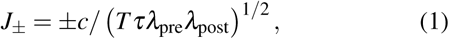

where *τ* is the timescale of synaptic impact, which is chosen by maximizing the model likelihood: *τ* = 0.004 s for the simulation data and *τ* = 0.001 s for the rat hippocampal data. The coefficient *c* is given as 5.16 for *p* = 0.001.

The connection parameter *J* is related to the post-synaptic potential (PSP), *w* mV. We approximate it with the linear relation

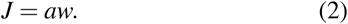

Here, the coefficient a is determined using synthetic data as *a* = 0.39 for the EPSP and *a* = 1.57 for IPSP. By combining this with equation (1), the necessary duration of spike recording needed to determine the likely presence of a connection of PSP is given as

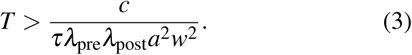

To make reliable inference, in addition to the above relation, it is also necessary to have collected a sufficiently large number of spikes during the interaction time window on the order of a few milliseconds. Here we require (METHODS):

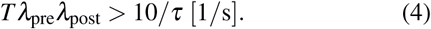

Table I shows the results of several cases of firing rates and the assumed PSPs using the *p* = 0.001. Unsurprisingly, to detect a weak connection for a low firing neuron requires gathering data for a long period of time. Figure 3A shows the connections estimated with different observation time windows, illustrating weak connections become visible as the recording duration increases.

**TABLE I.** Duration of spike recording required for verifying interneuronal connections, for the case of synaptic timescale *τ* = 0.001 s, which was selected for rat hippocampal data. The red letters represent the cases for which more than 2 h were required.

| Firing rates | EPSP: 5 mV | EPSP: 1 mV | EPSP: 0.5 mV | IPSP: 1 mV | IPSP: 0.5 mV |
| --- | --- | --- | --- | --- | --- |
| (10, 10) Hz | $T > 2$ min | $T > 30$ min | $T > 2$ h | $T > 2$ min | $T > 7$ min |
| (10, 5) Hz | $T > 3$ min | $T > 1$ h | $T > 4$ h | $T > 4$ min | $T > 10$ min |
| (5, 5) Hz | $T > 7$ min | $T > 2$ h | $T > 8$ h | $T > 7$ min | $T > 30$ min |
| (10, 1) Hz | $T > 20$ min | $T > 5$ h | $T > 20$ h | $T > 20$ min | $T > 1$ h |
| (5, 1) Hz | $T > 30$ min | $T > 10$ h | $T > 40$ h | $T > 40$ min | $T > 2$ h |
| (1, 1) Hz | $T > 3$ h | $T > 50$ h | $T > 200$ h | $T > 3$ h | $T > 10$ h |

**FIG. 3.**
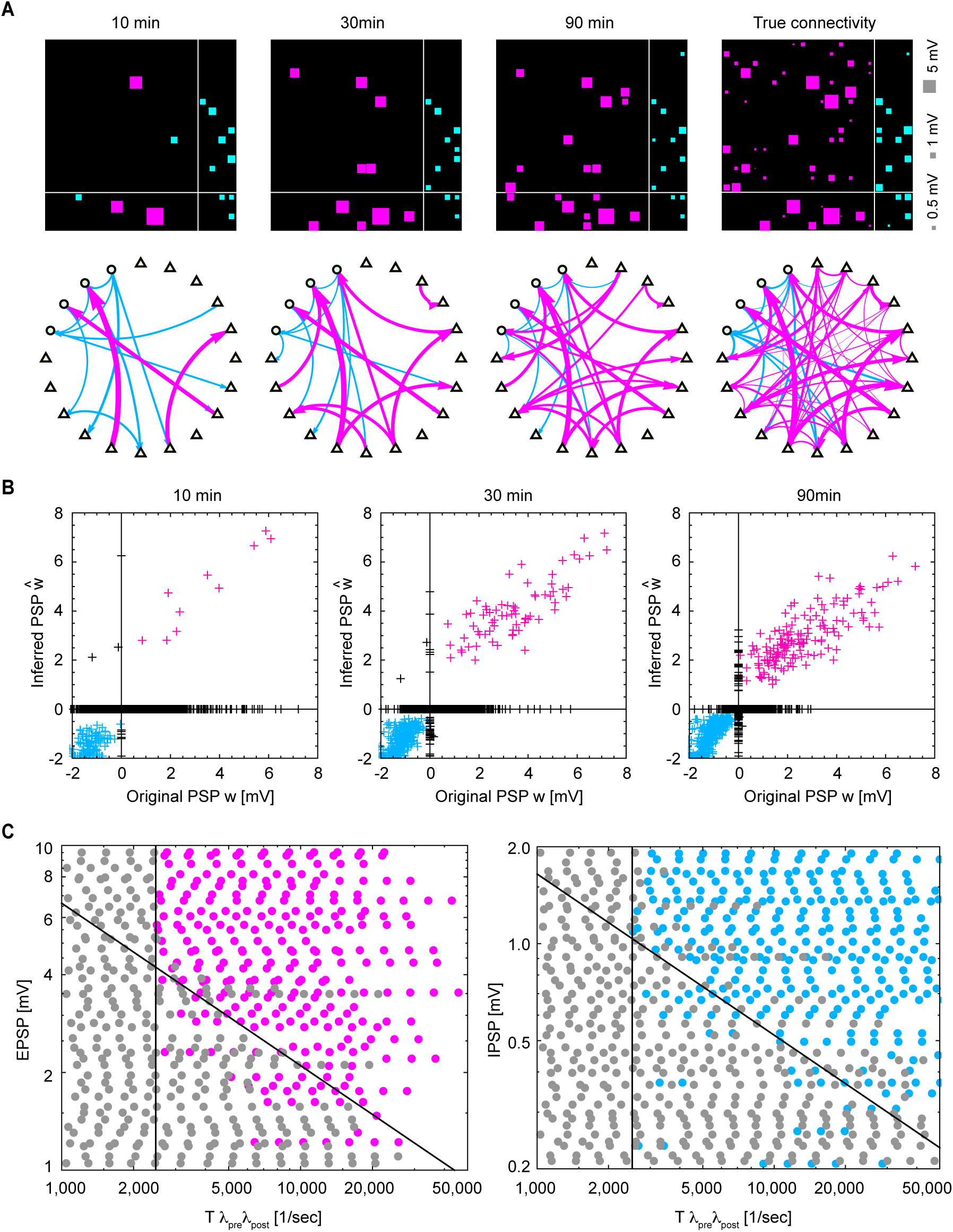
Neuronal circuits reconstructed from different observation time windows. (A) Interneuronal connections estimated from the observation time windows of 600, 1,800, and 5,400 s (10, 30, and 90 min) are plotted in reference to true connectivity. In each connection matrix, *x*-axis indicates reference neurons. In the network graphs shown in the second panel, excitatory and inhibitory neurons are depicted as triangles and circles, respectively. (B) Estimated postsynaptic potentials (PSPs) (*ŵ*) plotted against true parameters (*w*) were computed for 100 neurons randomly selected from the simulation. Points in the 1st and 3rd quadrants represent qualitatively correct inferences for excitatory and inhibitory connections (magenta and cyan, respectively). Points on the nonzero *y*-axis represent the false positive connections for unconnected pairs. Points on the nonzero *x*-axis represent the false negatives. (C) Detection status for connections of given PSPs with respect to the observation window (*T*). Connections estimated as excitatory and inhibitory are colored magenta and cyan, respectively, while undetermined ones are colored gray. Diagonal and vertical lines represent the theoretical formulas (3) and (4), respectively.

### D. Estimating PSPs

We believe that our method is of particular interest because it couches the connections in terms of PSPs for the individual neuronal pairs. Figure 3B compares the estimated PSPs (*ŵ*) against the true values (*w*) from the numerical simulation. Here we represent *ŵ* = 0 if the connection is undetermined, i.e., not significant. Thus, unconnected links (*w* = 0) that were classified as undetermined (true negatives) are placed at the origin. Points lying on the nonzero *x*-axis are existing connections that were not detected. Points lying on the nonzero *y*-axis are the functional or virtual connections that were estimated for unconnected pairs. The points in the 1st and 3rd quadrants represent true positives, or existing connections whose signs were correctly inferred as excitatory or inhibitory, respectively. The points in the 2nd and 4th quadrants are existing connections whose signs were misclassified.

The number of nonzero connections increases with the recording duration. Existing connections with large PSP amplitude tend to be detected with the signs correctly identified (points in the 1st and 3rd quadrants). There are also virtual connections assigned for unconnected pairs (nonzero *y*-axis). The number of such false positives is larger than the expected number of statistical errors (Figure 2A). This implies that the false connections may not be mere statistical fluctuations, but rather that they may reflect the functional connectivity indirectly connected via other unobserved neurons.

Figure 3C demonstrates the way individual connections emerge by increasing the recording duration. Here the abscissa is chosen as the observation window (*T*) multiplied by the firing rates of the preand postsynaptic neurons (λ_pre_ and λ _post_) so that all data are organized into a unified formula (inequality (3)). The values of *T*, λ _pre_, and λ _post_ for the excitatory connections tended to be smaller than those of inhibitory connections, because the firing rates of excitatory neurons were typically lower than those of inhibitory neurons.

### E. Excitatory*−*inhibitory (E*−*I) dominance index

The probability of misassigning individual connectivity for unconnected pairs was much higher than the statistical *p*value, because their firing is generally correlated with each other due to indirect interactions through unobserved neurons. Nevertheless, excitatory and inhibitory characteristics of individual neurons can be inferred with a lower error rate, because we can refer to multiple connections for each neuron.

We define an excitatory *−*inhibitory (E*−* I) dominance index as

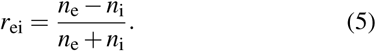

where *n*_e_ and *n* _i_ represent the numbers of identified excitatory and inhibitory connections innervated from each neuron, respectively. The E*−* I dominance indexes computed for 2 networks of 80 neurons each are plotted against firing rates of neurons (Figure 4A). In this case, excitatory and inhibitory characteristics of individual neurons were wellidentified based on E I dominance indexes. Inhibitory neurons typically exhibited higher firing rates in comparison to excitatory neurons. The firing irregularity measured using the local variation (*Lv*) of interspike intervals (Shinomoto, Shima, and Tanji, 2003; Mochizuki *et al.*, 2016) is plotted against firing rate. Spiking of inhibitory neurons tended to be more regular (smaller *Lv*) than that of excitatory neurons.

**FIG. 4.**
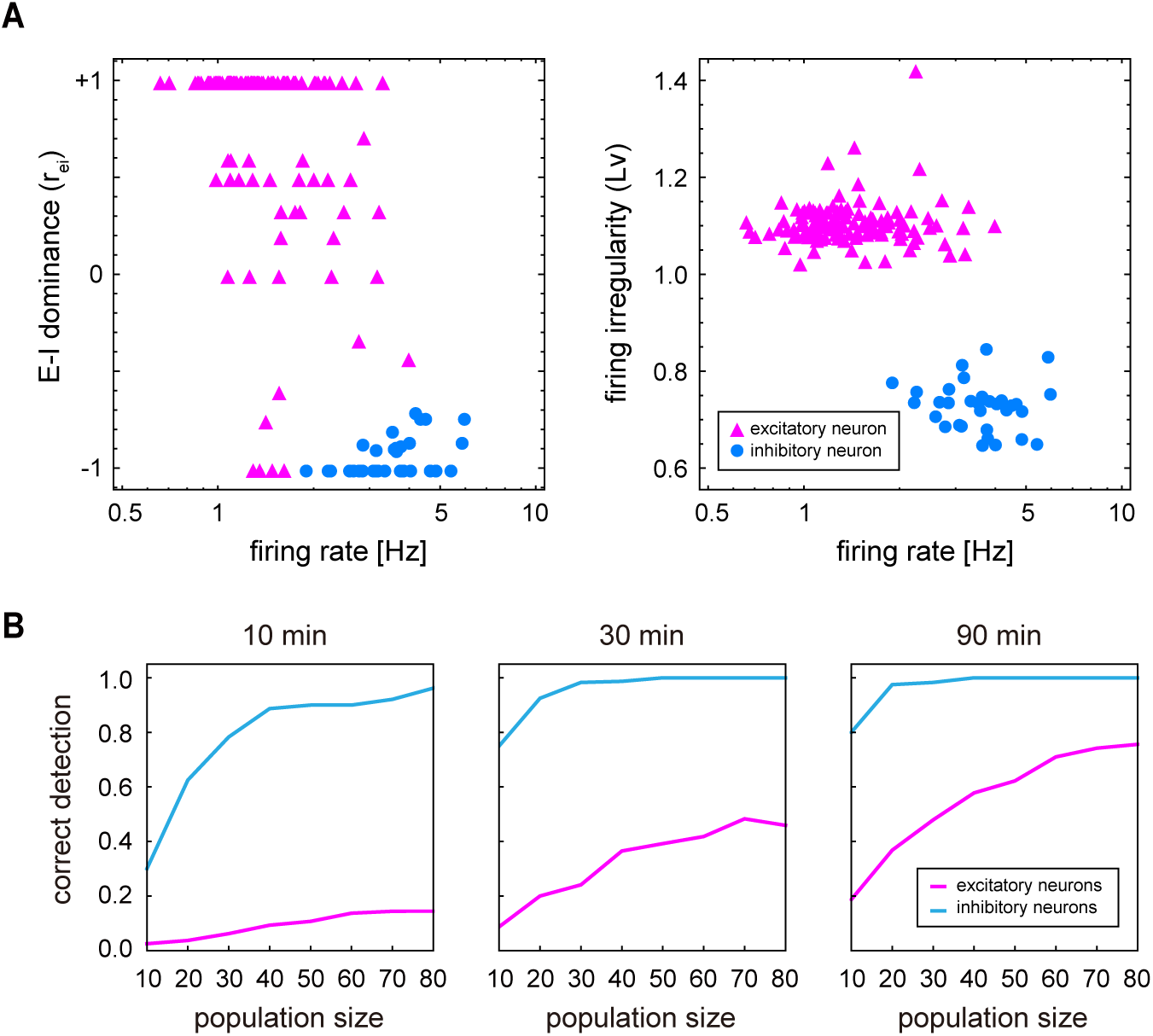
Excitatory− inhibitory (E− I) dominance index. (A) E − I dominance index *r*_ei_ = (*n*_e_ *n*_i_)*/*(*n*_e_ + *n*_i_) and the firing irregularity (*Lv*), plotted against the firing rate for 160 HH-type neurons (2 networks of 80 neurons). Excitatory and inhibitory neurons are plotted as triangles and disks, colored magenta and cyan, respectively. (B) The rates at which excitatory and inhibitory characteristics are identified correctly according to *r*_ei_ *>* 0 and *r*_ei_ *<* 0, respectively for excitatory and inhibitory neurons.

If we can record many spike trains in parallel for a long time, many excitatory and inhibitory neurons may be correctly identified according to *r*_ei_ *>* 0 and *r*_ei_ *<* 0, respectively. Figure 4B illustrates the manner in which the ratio of such correct identification depends on the total number of spike trains and the duration of observation.

### F. Real spike trains

We apply our method to spike trains recorded from the hippocampal CA1 area of a rat while it was exploring a square open field (Mizuseki *et al.* (2013): hc3 in CRCNS). Figure 5A displays the connections obtained with different observation time windows, demonstrating that more connections become visible as the recording duration increases, similar to the results seen with synthetic data. The connection matrix is divided into four quadrants according to the putative classification performed by manually analyzing waveforms, autocorrelograms, and mean firing rates (Skaggs *et al.*, 1996; Csicsvari *et al.*, 1998; Mizuseki *et al.*, 2009). We observe that connections in the 3rd, 4th, and 1st quadrants of the connectivity matrix representing excitatory-inhibitory, and inhibitory-inhibitory, and inhibitory-excitatory zones, respectively, are detected in a relatively short observation window. This is consistent with our formula (3) given that inhibitory neurons typically fire at high rates. Connections in the 2nd quadrant, representing the excitatory-excitatory zone, only appear after increasing the observation time window, and the estimated connection pattern remains sparse; more connections might have been identified if the observation period had been even longer. However, the estimated connection pattern is consistent with the finding using intracellular recording *in vitro* that inter-pyramidal connections in the hippocampus CA1 are sparse (Deuchars and Thomson, 1996).

**FIG. 5.**
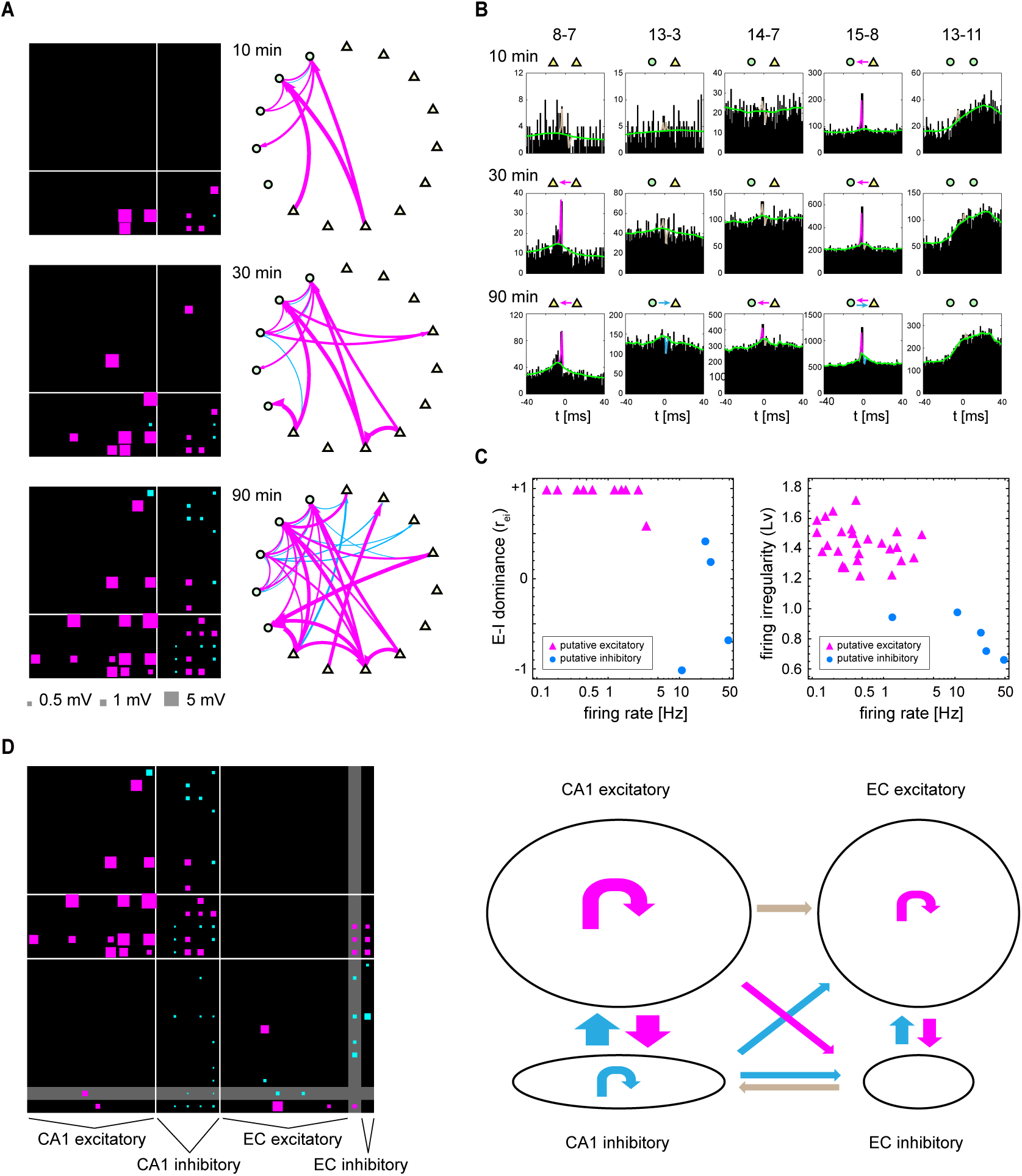
Neuronal circuits reconstructed from real spike trains *in vivo*. (A) Interneuronal connections estimated from spike trains recorded from the hippocampal CA1 area of a rat. Estimations were made with the observation time windows of 600, 1,800, and 5,400 s (10, 30, and 90 min). In each connection matrix, *x*-axis indicates reference neurons. The connection matrix is partitioned into groups of putative excitatory and inhibitory neurons defined manually according to other physiological cues such as waveforms. (B) Cross-correlations of several pairs of neurons computed at different time windows. The slow part of the GLM adapted to the data is depicted in green. The coupling filter is separately depicted in magenta, cyan, or gray, respectively, for the excitatory, inhibitory, or undetermined, respectively. Corroborated connections are indicated by arrows. (C) E − I dominance index (*r*_ei_) and the firing irregularity (*Lv*) plotted against the firing rates for putative excitatory and inhibitory neurons. (D) Estimated connections among neurons in CA1 and Entorhinal Cortex (EC). The connection matrix is partitioned into putative excitatory and inhibitory neurons in CA1 and EC. One EC unit, whose excitatory or inhibitory characteristic was not determined by the manual analysis is put in the gap (gray) between excitatory and inhibitory groups. In the network graph shown in the second panel, excitatory and inhibitory dominated connections are depicted in magenta and cyan, while connections of mixed characteristics are depicted in gray.

Figure 5B shows CCs of several neuron pairs. The crosscorrelations became less noisy as the observation time is increased, and some connections resolved (8-7, 13-3, 14-7, and 15-8). Some real spike trains exhibited large-scale wavy fluctuations (13-11), which may suggest that these neurons are under the influence of brain activity with lagged phases or perhaps they were responding to some unidentified external stimulus. Our method absorbs these fluctuations by adapting the slow part of the GLM (demonstrated as green lines), and succeeds in detecting a tiny synaptic impact by fitting coupling filters (lines colored magenta, cyan, and gray, respectively represent excitatory, inhibitory, and undetermined connections).

In Figure 5C, we plotted the E *−* I dominance index (*r*_ei_) and the firing irregularity (*Lv*) against the firing rate. The E− I dominance index is roughly consistent with the putative excitatory and inhibitory neurons. The irregularity of the putative excitatory neurons tended to be higher (larger *Lv*) than that of inhibitory neurons, similar to what we observed with the simulation data. The good separation of the putative excitatory and inhibitory neurons in these plots implies that we can classify recorded cells into excitatory and inhibitory neurons reliably without having to rely on their waveforms, because the E I dominance index, firing irregularity, and the firing rate are obtained solely from the spike times.

We also attempted to analyze a set of spike trains recorded simultaneously from multiple regions including CA1 and the Entorhinal Cortex (EC). Figure 5D demonstrates a matrix of estimated connections among excitatory and inhibitory neurons in CA1 and EC. Though the number of inter-regional connections was small in this sample data, our analysis method is generally applicable to any set of spike trains, irrespective of the recorded areas.

## III. DISCUSSION

We present a method for reconstructing neuronal circuitry from multichannel extracellular neuronal recordings. This method, based on a combination of the GLM and CC, can balance the antagonistic demands for reducing false positives and false negatives when estimating interneuronal connectivity. Our method is tolerant of the large variations in firing activity that often occur *in vivo*. As a critical part of the method, we show a framework for estimating the necessary duration of the spike recordings so that any likely interneuronal connections are detected. The duration is presented in terms of the firing rates of the preand postsynaptic neurons, and the presumed PSP.

It would be ideal to be able to estimate individual connections using intracellular or patch clamp recordings where the postsynaptic current caused by presynaptic neuronal firing can be measured, as is done with recordings from the rat cortex (Yoshimura, Dantzker, and Callaway, 2005; Song *et al.*, 2005; Buzsáki and Mizuseki, 2014). While those methods can reliably detect synaptic connections, they are limited because only a few neurons can be recorded simultaneously.

With the recent increase in parallel high channel count extracellular recordings from anaesthetized and behaving animal subjects (Buzsáki, 2004; Jun *et al.*, 2017), it is possible to estimate the connection strength between a number of neurons (Fujisawa *et al.*, 2008; Bock *et al.*, 2011; Gong *et al.*, 2015). Several strong analytical methods for estimating connections from spike trains have been developed, including the cross-correlation analysis (Perkel, Gerstein, and Moore, 1967; Toyama, Kimura, and Tanaka, 1981; Sakurai, 1996; Ventura, Cai, and Kass, 2005; Doiron *et al.*, 2016) and the GLM (Okatan, Wilson, and Brown, 2005; Truccolo *et al.*, 2005; Pillow *et al.*, 2008; Chen *et al.*,2011;Kim *et al.*,2011;Kobayashi and Kitano,2013;Zaytsev, Morrison, and Deger, 2015; Gerhard, Deger, and Truccolo, 2017). While CCs have been used to estimate neuronal connectivity, the classical cross-correlation analysis becomes unreliable when there are large fluctuations in the data. One approach to solving this problem has been to jitter the time stamps of spikes (Amarasingham *et al.*, 2012; Schwindel *et al.*, 2014). Another approach has been to apply GLM to parallel spike trains. However, the size of the computation increases as the recording time increases. Because the number of neuronal pairs increases by the square of the number of spike trains (e.g., 10,000 pairs should be examined for 100 parallel spike trains), computation for estimating individual connections of each pair should be modest. Our analysis can be conducted with a reasonable computation time with amounts of data that can reasonably be collected, because our GLM analyzes the CC for a time window of 100 ms rather than the entire spike trains. Our GLM may also adapt to wavy fluctuations in CC, making it tolerant to large-scale fluctuations that are often attendant on real spike trains *in vivo*.

Because recording time is limited, a possible restriction on inferring connectivity might be that there is not enough data. Even if a given neuron fired several times with each spike occurring shortly after the firing of an index neuron, there still might not be enough data to conclude that there is a synaptic connection. Here we made estimates on the duration of spike recordings needed so that any likely interneuronal connections will be detected (cf. Table I).

When we applied our method to data recorded from the rat hippocampus we identified connections for four types of pairs, excitatory-excitatory, excitatory-inhibitory, inhibitory-inhibitory, and inhibitory-excitatory, in numbers consistent with those identified physiologically (Mizuseki *et al.*, 2009), supporting the efficacy of our method. Typically, the pyramidal neurons have low background firing rates and interneurons have higher firing rates. Our analysis (cf. inequality (3)) indicates that the necessary recording duration is inversely proportional to the product of the firing rates of the preand postsynaptic neurons. Thus, connections between neurons firing at high frequencies can be detected with a relatively short observation duration. For neurons with low firing rates, on the other hand, data will have to be collected for much longer periods, and we expect that excitatory-excitatory connections will be detected only if there is a relatively long recording period. The consequences of this have been seen with experimental data; for instance, synapses that connect with inhibitory interneurons were frequently detected, and connections between excitatory neurons were rarely detected (Barthó *et al.*, 2004; Fujisawa *et al.*, 2008). The hippocampal data analyzed in this study (Figure 5A) conforms to this pattern, and our analysis provides insight into how this happens.

Our approach and method provide a means for estimating a map of neuronal connections from high channel count simultaneous recordings. We presume, based on anatomical differences, that these maps will have different structures in different functional brain regions. Having a reliable technique for estimating the maps offers the opportunity to identify these different structures, thereby providing a basis for understanding the differences in information processing that arises from differences in anatomy and connected structures.

## IV. METHODS

### A. Estimating interneuronal connectivity

Here we describe our GLM analysis, the bases of validating connections and selecting a *p* value, and the method of estimating the post-synaptic potential (PSP).

#### CC-GLM

To discover interneuronal connections between a pair of neurons, we devise a GLM that detects short-term synaptic impacts in the CC (as schematically depicted in Figure 1A and as real cross-correlograms of rat hippocampal data in Figure 5B). We design the GLM describing the CC (which we call CC-GLM) as

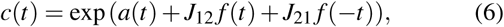

where *t* is the time from the spikes of the reference neuron, and *a*(*t*) represents large-scale fluctuations produced outside the pair of neurons. *J*_*i* *j*_ represents interneuronal connection from the *j*th neuron to the *i*th neuron. The time profile of the synaptic interaction is modeled as 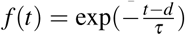 for *t > d* and *f* (*t*) = 0 otherwise, where *τ* is the typical timescale of synaptic impact and *d* is the transmission delay. The connection parameter *J*_*ij*_ of our CC-GLM can be derived from a model of the original interaction process between neurons (Supplementary Information A).

Given an underlying rate *c*(*t*), the probability for spikes to occur at {*t*_*k*_} = {*t*_1_,*t*_2_, *·,t*_*N*_} is obtained theoretically as Cox and Lewis (1966); Daley and Vere-Jones (2003),

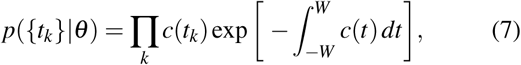

where θ represents a set of parameters that characterize *c*(*t*).

To detect short-term synaptic impacts of a few ms hidden in large-scale fluctuations, we make *a*(*t*) adapt to slow variation in the CC. We implement this by discretizing the temporal modulation *a*(*t*) at intervals of Δ*t* = 1 ms in a time window of 2*W* = 100 ms, which is represented by a set of parameters 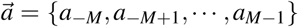. We assume that the background activity is slowly modulated. This may be implemented by providing a prior distribution that penalizes a large gradient or (*da/dt*) ^2^. In terms of discretized parameters, we give the prior distribution:

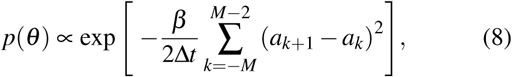

where β is a hyperparameter representing the flatness of *a*(*t*); *a*(*t*) is nearly constant if β is large, or is rapidly fluctuating otherwise. Here, the hyperparameter β is selected as β = 1000 ms. For the connection parameters *J*_12_ and *J*_21_, we assume uniform priors.

The posterior distribution of a set of parameters 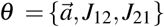, given the spike data {*t*_*k*_}, is obtained from Bayes’rule as

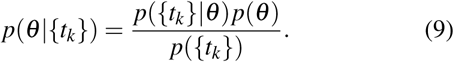

The parameters are determined with the maximum *a posteriori* (MAP) estimate, that is, by maximizing the posterior distribution or its logarithm:

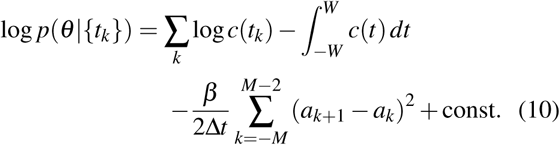

The maximization is performed efficiently using the Levenberg–Marquardt method (Supplementary Information B).

#### Statistical test for determining connectivity

We determine the presence of an interneuronal connection by disproving the null hypothesis that a connection is absent. Namely, we conclude that a connection is likely present if the estimated parameter is outside the confidence interval for the null hypothesis; otherwise, the presence of a connection is undetermined. The null hypothesis is that two neurons generate spikes at their baseline firing rates independently of each other. According to Poisson statistics, the variance of the number of spikes generated in a time interval Δ after the spike of a reference neuron is equal to its mean. The mean spike number is obtained by multiplying the intensity *c*(0) by an interval Δ,

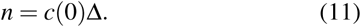

Assuming that the connection *J* is small, the average number of spikes caused by an interneuronal connection during an interval Δ is approximated as

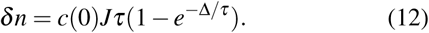

The condition that the synaptic interaction produces a significant impact in the CC is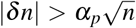, where α_*p*_ is the alpha-value representing a threshold for the normal distribution (α_*p*_ = 2.58 for *p* = 0.01 and α_*p*_ = 3.29 for *p* = 0.001). In terms of the estimated connection parameterĴ, this conditionis given as

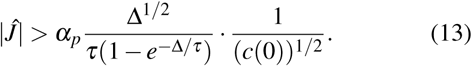

Here, Δ^1/2^/(*τ*(1 − *e−*Δ*/τ*)) in the right-hand-side of this inequality is bounded from below by 1.57*τ−*1*/*2 (Δ = 1.26*τ*). Thus we have the following inequality:

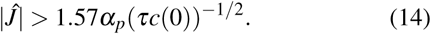

The typical duration of spike recording needed for the connectivity inference (inequality (3)) is obtained from equation (14) by approximating *c*(0) = *T* λ_pre_λ_post_, where *T* is the total duration of recording.

Another requirement is that spike trains should contain a sufficiently large number of spikes to make a reliable inference. A typical number of spikes contained in the CC in the interaction time window is *T* λ_pre_λ_post_*τ*. By requiring this to be greater than 10, we obtain the inequality (4).

#### Selecting the *p* value

Although we obtained the confidence interval of the connection parameter *J*_*i* *j*_ at a given *p* value above, the probability of assigning spurious connectivity to anatomically disconnected pairs is higher than the given *p* value, because spike trains are correlated. Such spurious connections or falsepositives may be reduced by decreasing the *p* value. However, this operation may cause the vast majority of existing connections to be missed, thus producing a huge number of falsenegatives. Thus, the *p* value should be chosen so that these conflicting demands (of reducing false positives and false negatives) are optimally compromised.

As we can directly count false positives and false negatives in simulation data, we may select a *p* value such that the performance of the inference is maximized. As a measure for assessing the performance of connectivity inference, we adopt the Matthews correlation coefficient (MCC) (Matthews, 1975) defined as

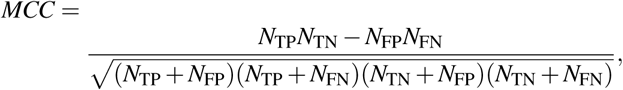

where *N*_TP_, *N*_TN_, *N*_FP_, and *N*_FN_ represent the numbers of true positive, true negative, false positive, and false negative connections, respectively.

Because there are excitatory and inhibitory connections, we may obtain two coefficients for individual categories. To evaluate the quality of inference in terms of a single measure, here we take the macro-average MCC that sets equal importance to these categories (Yang, 1999; Sun and Lim, 2001):

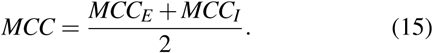

In computing the coefficient for the excitatory category *MCC*_*E*_, we classify connections as excitatory or other (disconnected and inhibitory); for the inhibitory category *MCC*_*I*_, we classify connections as inhibitory or other (disconnected and excitatory).

#### Estimating PSPs from GLM connection parameters

We translate the GLM connection parameters *J*_*i* *j*_ into biological PSPs *w*_*i* *j*_ mV. This relation is obtained by numerically simulating a network of neurons interacting through known connections {*w*_*i* *j*_ }and by applying the GLM to their spike trains to estimate the connection parameters *J*_*i* *j*_. Regarding synaptic connections *w*_*i* *j*_ for which {*J*_*i* *j*_ *}*was verified in the correct signs, we assume a linear relation as in equation (2):

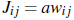

The coefficients *a* is determined by applying regression analysis to the data. We obtained *a* = 0.39 for EPSP and 1.57 for IPSP, respectively.

When we newly estimate connection parameters 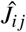 from spike trains, they can be translated into PSPs using the relation:

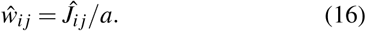

Figure 3B compares the estimated PSPs *ŵ*_*ij*_ with the original PSPs values *w*_*i* *j*_ of a model neural network.

In our numerical simulation, synaptic connectivity is given in terms of conductance. Thus we have to translate conductance into PSP. The translation rule is described in Supplementary Information C.

### B. Synthetic data

We ran a numerical simulation of a network of 1,000 Hodgkin–Huxley (HH) type neurons interacting through fixed synapses. Of them, 800 excitatory neurons innervate to 12.5 % of other neurons with EPSPs that are log-normally distributed (Song *et al.*, 2005; Teramae, Tsubo, and Fukai, 2012; Buzsáki and Mizuseki, 2014), whereas 200 inhibitory neurons innervate randomly to 25 % of other neurons with IPSPs that are normally distributed.

#### Neuron models

For excitatory pyramidal cells, we adopted HH type models developed by Destexhe and Paré (1999). The membrane potential *V* obeys the equation:

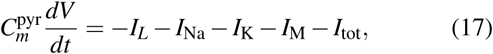

where 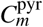 is the membrane capacitance,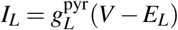 is the leak current,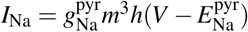 is the Na+ current,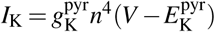 is the delayed-rectifier K+ current,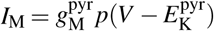 is the muscarinic potassium current, and *I* tot is the total input current from the other neurons. The gating variables *x ∈ {m, h, n, p}* are described by the kinetic equation:

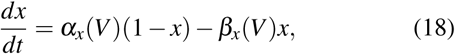

where α_*x*_ and β_*x*_ are the activation and inactivation functions, respectively. The activation and inactivation functions and the parameter values are summarized in Table II.

**TABLE II.** Parameters for pyramidal neurons and interneurons

| Neuron Models |  |
| --- | --- |
| $C_m^{\text{pyr}}, C_m^{\text{inh}} [\mu\text{F}/\text{cm}^2]$ | 1.0, 1.0 |
| $g_L^{\text{pyr}}, g_L^{\text{inh}} [\text{mS}/\text{cm}^2]$ | 0.045, 0.1 |
| $g_{\text{Na}}^{\text{pyr}}, g_{\text{Na}}^{\text{inh}} [\text{mS}/\text{cm}^2]$ | 50.0, 112.0 |
| $g_K^{\text{pyr}}, g_M^{\text{pyr}}, g_{\text{Kv}3}^{\text{inh}}, g_{\text{Kv}1}^{\text{inh}} [\text{mS}/\text{cm}^2]$ | 5.0, 0.07, 224.0, 0.224 |
| $E_L^{\text{pyr}}, E_L^{\text{inh}} [\text{mV}]$ | -80.0, -70.0 |
| $E_{\text{Na}}^{\text{pyr}}, E_{\text{Na}}^{\text{inh}} [\text{mV}]$ | 50.0, 55.0 |
| $E_K^{\text{pyr}}, E_K^{\text{inh}} [\text{mV}]$ | -90.0, -97.0 |
| $S [\text{cm}^2]$ | $3.5 \times 10^{-4}$ |

For inhibitory interneurons, we adopted HH type models developed by Erisir *et al.* (1999). The membrane potential *V* obeys the equation:

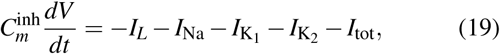

where 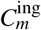 is the membrane capacitance,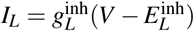 is the leak current,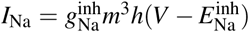 is the Na+ current,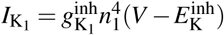 and 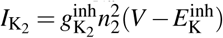 are the delayed-rectifier K+ current due to Kv1.3 and Kv3.1-Kv3.2 conductance, respectively, and *I*_tot_ is the total input current. The gating variables *x ∈* {*m, h, n*_1_, *n*_2_ }follow the kinetic equation (18), with the activation and inactivation functions prescribed by the original paper (Erisir *et al.*, 1999). The parameter values are summarized in Table II.

#### Synaptic connections

Each neuron receives synaptic currents induced by the firing of other neurons. Excitatory synaptic currents are mediated by 2-amino-3-(5-methyl-3-oxo-1,2-oxazol-4-yl) propanoic acid (AMPA) and N-methyl-D-aspartate (NMDA) receptors, whereas inhibitory synaptic currents are mediated by γ-aminobutyric acid (GABA)-A receptors. The total input current to the *i*th neuron is given by

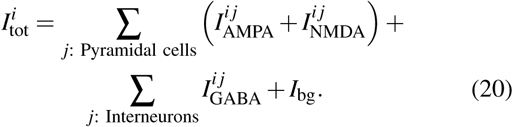

where 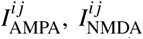,and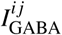, respectively represent the synaptic currents given by the AMPA, NMDA, and GABA receptors, and *I*_bg_ represents the background current.

For AMPA-mediated current, we adopted the depressing synapse model proposed by Tsodyks and Markram (1997)

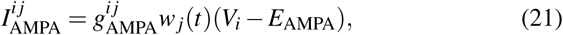

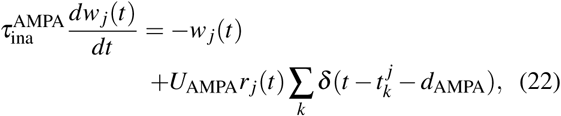

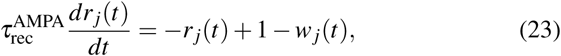

where 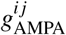 is the maximal AMPA conductance, *Vi* is the membrane potential of the postsynaptic neuron,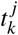 is the *k*th spike time of the pre-synaptic neuron, and *d*_AMPA_ is the synap-tic conduction delay. For each connection, the conduction delay is drawn from a uniform distribution between 0 and 2 ms. *w*_*j*_ and *r* _*j*_ represent the fraction of synaptic resources in the effective and recovered states, respectively. The AMPA parameter values are summarized in Table III.

**TABLE III.**
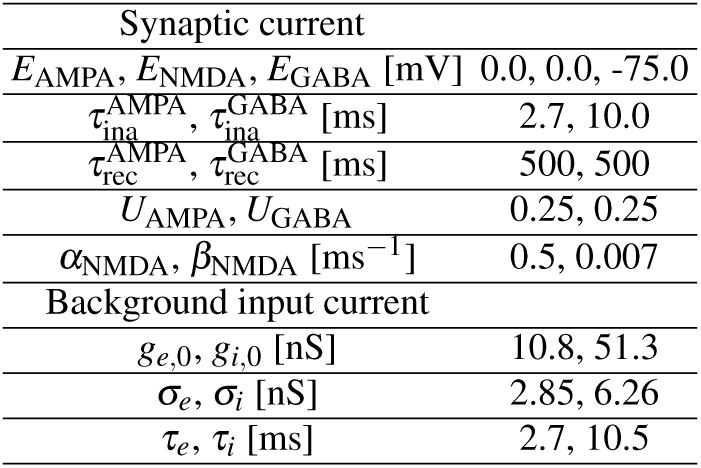
Parameters for synaptic currents and background inputs.

For NMDA-mediated current, we adopted the first-order kinetic equation proposed by Destexhe, Mainen, and Sejnowski (1998)

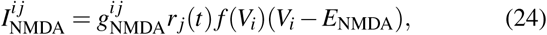

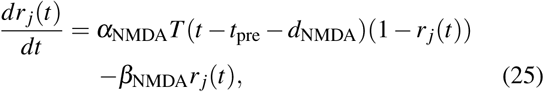

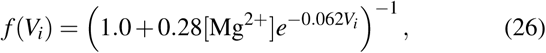

where [Mg2+]= 1.0 mM is the extracellular magnesium concentration, *t*_pre_ is the last spike time of the presynaptic neuron, *d*_NMDA_ is the conduction delay drawn from a uniform distribution between 0 and 2 ms, and *T* (*t*) represents the transmitter concentration in the cleft. When a spike occurs in a pre-synaptic neuron, a transmitter pulse is induced such that *T* (*t*) = 1 mM for a short period (1 ms) and the concentration returns to *T* (*t*) = 0. The NMDA parameter values are summarized in Table III.

For GABA-A-mediated current, we adopted the depressing synapse model proposed by Tsodyks and Markram (1997)

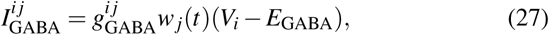

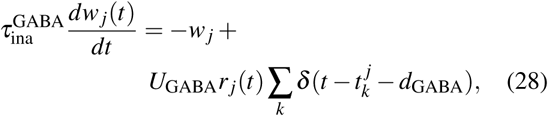

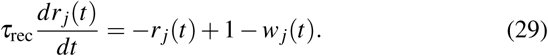

where *d*_GABA_ is the conduction delay drawn from a uniform distribution between 1 and 3 ms. The GABA parameter values are summarized in Table III.

We ran a simulation of a network consisting of 800 pyramidal neurons and 200 interneurons interconnected with a fixed strength. Each neuron receives 100 excitatory inputs randomly selected from 800 pyramidal neurons and 50 inhibitory inputs selected from 200 interneurons.

The AMPA conductance 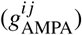 is drawn independently from a log-normal distribution (Song *et al.*, 2005; Teramae, Tsubo, and Fukai, 2012)

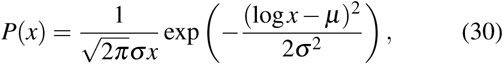

where *μ* =-3.37 and σ = 1.3 are the mean and SD of the natural logarithm of the AMPA conductance. The NMDA and GABA conductances (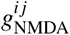 and 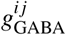) are sampled from the normal distribution

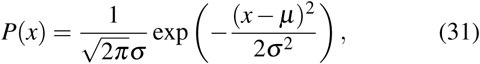

where *μ* and σ are the mean and SD of the conductances. Parameters are *μ*_NMDA_ = 8.5×10^−4^ mS/cm^2^, σ_NMDA_ = 8.5×10^−5^ mS/cm^2^ and *μ*_GABA_ = 0.34 mS/cm^2^, σ_GABA_ = 0.27 mS/cm^2^ for the NMDA and GABA conductance, respectively. If the sampled value is less than zero, the conductance is resampled from the same distribution.

Because our model network is smaller than real cortical networks, where each neuron receives inputs from the order of 10,000 neurons (Braitenberg and Schüz, 1998), we added a background current to represent inputs from many neurons, as previously done by (Destexhe *et al.*, 2001). The background current is given as the sum of excitatory and inhibitory inputs:

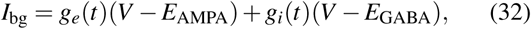

where the total excitatory and inhibitory conductance *g*_*e,i*_(*t*) obey the Ornstein–Uhlenbeck process (Tuckwell, 1988), representing random bombardments from a number of neurons.

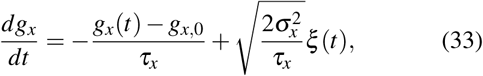

where *x* represents excitatory (*e*) or inhibitory (*i*), *g*_*x*_ and σ_*x*_ are the mean and SD of the conductance, *τ*_*x*_ is the synaptic time constant, and *ξ*(*t*) is the Gaussian white noise with zero mean and unit variance. Parameters for the background inputs are summarized in Table III.

#### Numerical simulation

Simulation codes were written in C++ and parallelized with MPI programming. Simulations were conducted on a computer cluster with 64 nodes, each consisting of 4 processors. The time step was 0.01 ms for excitatory (pyramidal) neurons and 0.001 ms for inhibitory (inter) neurons. The neural activity was simulated up to 10,000 s. In practice, it took about two weeks to perform the entire simulation. Simulation codes are available upon request.

### C. Experimental data

Spike trains were recorded from the hippocampal area of a rat, while it was exploring an open square field. Experimental procedures, data collection, and spike sorting are as described in detail in Mizuseki *et al.* (2009). All protocols were approved by the institutional animal care and use committees of Rutgers University and New York University. Hippocampal principal cells and interneurons were separated on the basis of their waveforms, auto-correlograms, and mean firing rates (Skaggs *et al.*, 1996; Csicsvari *et al.*, 1998; Mizuseki *et al.*, 2009). All data used in this paper can be found in hc3 data sets at CRCNS (Mizuseki *et al.* (2013); CRCNS.org. http://dx.doi.org/10.6080/K09G5JRZ).

## ACKNOWLEDGMENTS

This study was supported in part by JSPS KAKENHI Grant Number 17H03279 and 18K11560, JST ACT-I Grant Number JPMJPR16UC, the Okawa Foundation for Information and Telecommunications, and the open collaborative research and MOU grant at the National Institute of Informatics in Japan to R.K., and JSPS KAKENHI Grant numbers 26280007 and 17H06028 and JST CREST Grant Number JPMJCR1304 to S.S. We thank Yuzuru Yamanaka, Tatsuya Goto, and Kazuki Fujita for constructive comments on this manuscript.

